# The cell wall regulates dynamics and size of plasma-membrane nanodomains in *Arabidopsis*

**DOI:** 10.1101/455782

**Authors:** JF McKenna, DJ Rolfe, SED Webb, AF Tolmie, SW Botchway, ML Martin-Fernandez, C Hawes, J Runions

**Affiliations:** Department of Biological and Medical Sciences, Oxford Brookes University, Sinclair Annex, Gipsy Lane, OX3 0BP; Central Laser Facility, Research Complex at Harwell, Science and Technology Facilities Council, Rutherford Appleton Laboratory, Oxfordshire OX11 0QX, United Kingdom; Present address: BBSRC (UKRI), Polaris House, N Star Ave, Swindon, SN2 1UH, United Kingdom

## Abstract

Plant plasma-membrane (PM) proteins are involved in several vital processes, such as detection of pathogens, solute transport and cellular signalling. Recent models suggest that for these proteins to function effectively there needs to be structure within the PM allowing, for example, proteins in the same signalling cascade to be spatially organized. Here we demonstrate that several proteins with divergent functions are located in clusters of differing size in the membrane using sub-diffraction-limited Airyscan confocal microscopy. In addition, single particle tracking reveals that these proteins move at different rates within the membrane. We show that the actin and microtubule cytoskeletons appear to significantly regulate the mobility of one of these proteins (the pathogen receptor FLS2) and we further demonstrate that the cell wall is critical for the regulation of cluster size by affecting single particle dynamics of two proteins with key roles in morphogenesis (PIN3) and pathogen perception (FLS2). We propose a model in which the cell wall and cytoskeleton are pivotal for differentially regulating protein cluster size and dynamics thereby contributing to the formation and functionality of membrane nanodomains.

**One sentence summary:** Size and mobility of protein nanodomains in the plant plasma-membrane are regulated by interaction with the cell wall extracellular matrix.

**Significance statement:** The plant plasma membrane acts as the front line for cellular perception of the environment. As such, a large number of signalling and transport proteins which perceive or transport environmental signals, developmental cues and nutrients are located within it. Recently, a number of studies have revealed that proteins located within the plasma membrane do not simply freely diffuse within its plane. Rather, proteins are localized in nanometer sized structures called nanodomains. In addition to the plasma-membrane, plant cells also have an extracellular matrix, the cell wall. Here we have shown that the cell wall has a role in regulating the dynamics and size of plasma membrane nanodomains for proteins involved in morphogenesis (PIN3) and pathogen perception (FLS2).

## Introduction

The plasma membrane (PM) plays key roles in compartmentalization and protection of cells from the environment (1). In plants, proteins located within the PM are critical for signal perception, transduction and the controlled import and export of molecules (2). The structure of the PM was described by the fluid mosaic-model as a diffuse mixture of proteins in motion (3). However, this does not fit observations of protein spatial heterogeneity in membranes and subsequent models have been developed (4) which incorporate lipid rafts, detergent resistant membrane regions, cytoskeleton corralling and extracellular matrices as mechanisms of spatial constraint (5).

While proposed models of PM organization are under dispute and no single model explains all experimental observations across different model organisms, a number of proteins are known to locate to specific domains in the plant PM. The best studied of these in plants is the REMORIN family (6–8). Members of the REMORIN family form non-overlapping PM nanodomains (6). We define nanodomains here as others have previously: distinguishable submicron protein or lipid assemblies which are 20nm to 1μm in size (8). While the patterning of these REMORIN nanodomains has been well described, only recently has a molecular function of these proteins been demonstrated. In rice, OsREM4.1 is upregulated by abscisic acid and interacts with OsSERK1 to downregulate brassinosteroid signalling (9). Additionally, *Medicago* SYMREM1 is a key protein involved in segregating the receptor LYK3 into stable nanodomains and functioning during host cell infection (10). Proteins critical for normal morphogenesis and development such as PIN1 and PIN2 are localized to defined domains in the PM. PIN2 has been shown, using STED super-resolution imaging, to form clusters in the PM, with controlled endo-, and exocytosis from adjacent membrane regions to the localization domain (11). Additionally, the pathogen receptor FLS2 has been shown to localize to nanodomains in the plasma-membrane (12). Spatial organization of proteins in the PM is, therefore, important for development and response to the environment, but how is membrane domain patterning regulated?

The underlying cytoskeleton and outlying cell wall can be thought of as a continuum with the PM (2, 13). There are numerous examples of cytoskeletal and PM mechanisms which play roles in cell wall production and regulation of cell wall patterning: i) the microtubule-guided CesA complex determines patterns of cellulose microfibril deposition (14, 15), ii) microtubule-associated MIDD1 is involved in secondary cell-wall pit formation (16), iii) the CASP family of proteins form a PM nanodomain which defines the site of Casparian strip formation (17), and iv) FORMIN1 is anchored within the cell wall, spans the PM and nucleates actin filaments as part of a mechanism in which cell-wall anchoring is required for actin cytoskeleton organisation (18). The cell wall has been shown to have a role in regulating the lateral diffusion of two ‘minimal’ membrane proteins which have GFP projecting into the cell wall space (5). ‘Minimal’ membrane proteins are artificially-created peptides which localise to the plasma membrane via one of a number of association mechanisms. They were designed as fluorescent protein fusions and have no predicted protein interactions or biological functions. The plant cell wall is also required for normal localisation of PIN2 in the membrane and hence regulation of cell polarity (19). These examples highlight the possibility that the components of the cytoskeleton / PM / cell wall continuum can regulate each other, with cell-wall regulation of plasma-membrane, and cytoskeleton organization already observed (5, 20).

A systematic study of a number of PM proteins in transiently and stably expressing plant cells has demonstrated a difference in their lateral mobility (5). This was achieved by Fluorescence Recovery After Photobleaching (FRAP) using high temporal but low spatial resolution. An ever increasing toolkit of sub-diffraction limited microscopy techniques has been developed over recent years and we have used Airyscan imaging (21, 22) of flat membrane sheets in *Arabidopsis thaliana* hypocotyl cells to image PM structure with high spatial resolution. We chose to use Airyscan imaging and Total Internal Reflection Fluorescence - Single particle (TIRF-SP) imaging as they do not involve the use of special fluorophores required for PALM or a high power depletion laser used in STED which causes damage of aerial tissue in plants due to the presence of light absorbing chloroplasts. A combination of TIRF-SP and Airyscan imaging allows fast temporal acquisition with sub-diffraction limited resolution (down to 140 nm for the latter) in all plant tissues with the use of any existing fluorophore (21).

We show that FLS2, PIN3, BRI1 and PIP2A, form clusters of differing size from 164 to 231 nm. Our investigation indicates that actin and microtubule cytoskeletons regulate the diffusion rate of the pathogen receptor FLS2 but not the hormone transporter PIN3. Furthermore, cluster size and diffusion rate of both FLS2 and PIN3 are regulated by cellulose and pectin components of the cell wall.

We hypothesise that the constraint of the cell wall on PM proteins and differential regulation by the actin and microtubule cytoskeletons can contribute to PM organisation by altering protein dynamics and hence nanodomain size. This is a mechanism by which proteins can exist within different sized nanodomains.

## Results

### Plasma-membrane proteins form clusters within the membrane

We chose to study several well characterized PM proteins which have a variety of functions in order to determine how different proteins are organized in the PM and whether their dynamic behaviors differ. Airyscan imaging and determination of nanodomain full width half maximum (FWHM) demonstrated that proteins form clusters within the PM which are not resolved by diffraction-limited confocal imaging (Fig.1. & S1). Protein clusters were observed and measured for the auxin transporter PIN3 (Puncta FWHM, = 166.7±31.1 nm, Fig.1), the pathogen receptor FLS2 (Puncta FWHM = 164.3±32.0 nm, Fig.1), the hormone receptor BRI1 (Puncta FWHM = 172.6±41.3 nm, Fig.S1) and the aquaporin PIP2A (Puncta FWHM = 194.3±66.8 nm, Fig.S1). Cluster diameter was determined by FWHM measurements of line profiles over randomly selected nanodomains. Each protein observed had a nanodomain diameter below the theoretical 250nm Abbe resolution limit of confocal microscopy using GFP (Fig.1D)(23). When compared to REM1.3 (Puncta FWHM = 231.0±44.8 nm, Fig.1) which is known to form highly stable nanodomains resolvable by confocal microscopy within the PM (6), FLS2 and PIN3 clusters are significantly smaller and are more dynamic within the membrane (Fig.1C and S1).

**Figure 1.**
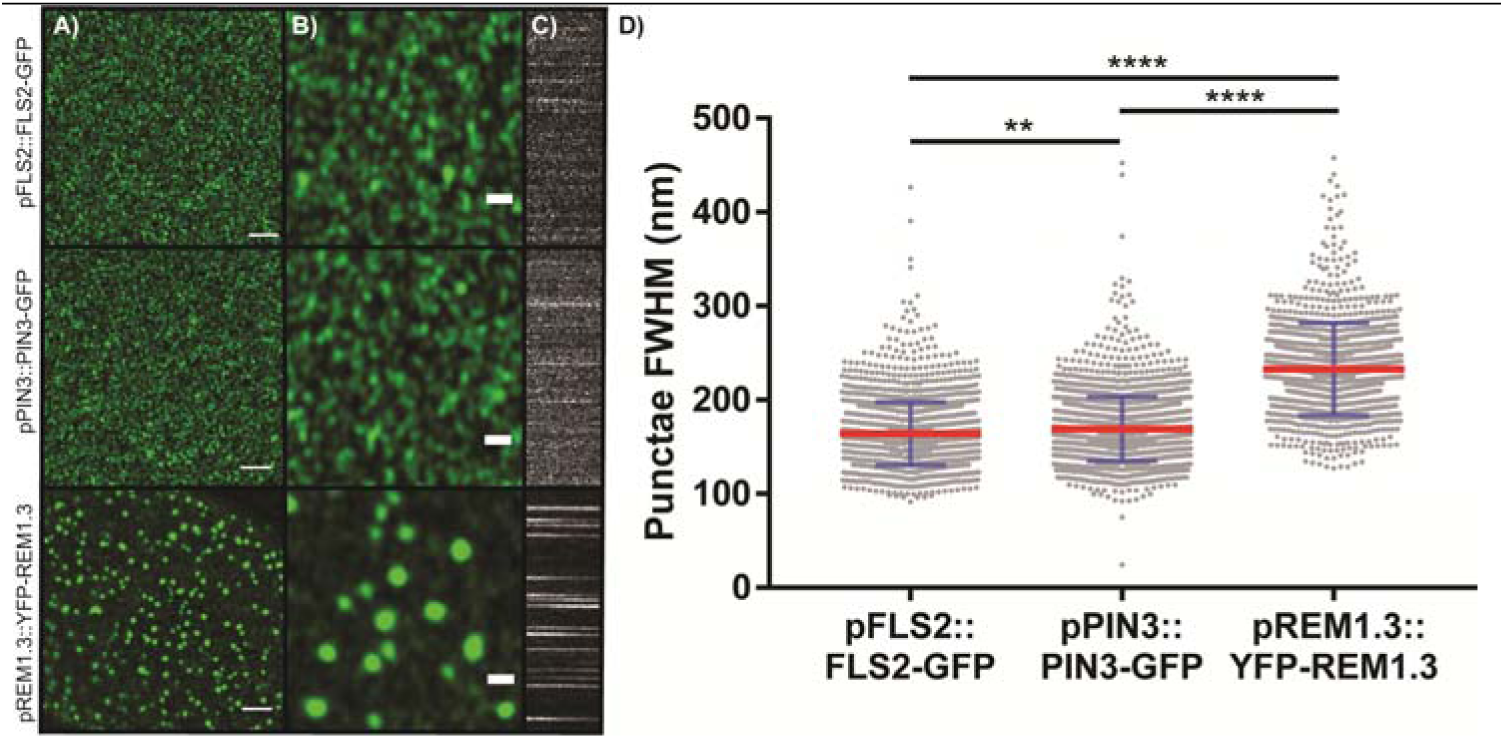
PM proteins form clusters in the hypocotyl membrane. A) Airyscan imaging of pFLS2::FLS2-GFP, pPIN3::PIN3-GFP and pREM1.3::YFP-REM1.3 clusters in the membrane of stably-transformed *A. thaliana*, scale bar = 2μm. B) Digitally magnified image of those in A) showing clusters in more detail, scale bar= 500nm. C) Kymographs showing dynamics of each nanocluster in A) over time where x = time, y =line profile. D) Box-and-whisker plot of full width half maximum (FWHM) measurement of cluster diameter for PM proteins in A). Red line indicates mean, blue error bars represent standard deviation. Nanodomain diameter differs significantly for each protein pair. **=p <0.01 and ****=p <0.0001, ANOVA with multiple comparisons.

### Proteins move at different speeds within the membrane

To determine the diffusion rate of select proteins within the PM we used Total Internal Reflection Fluorescence - Single Particle Tracking (TIRF-SPT) which yields high spatial and temporal resolution tracking information. We chose to focus on the PM proteins p35S::paGFP-LTI6b, p35S::PIP2A-paGFP, pFLS2::FLS2-GFP and pPIN3::PIN3-GFP as these cover a diverse range of functions from pathogen perception, to morphogen transport and resource acquisition (Fig.2, Supplemental movie 1). It is worth noting, TIRF-SP imaging and tracking can be performed with both photoactivatable GFP (paGFP) and GFP with overexpression or native promoters. However, expression needs to be within a range sufficient for signal detection but not so bright as to saturate the detector. This was the case for GFP-linked protein expression driven by the PIN3 and FLS2 promoters in the *A. thaliana* hypocotyl. Here we show diffusion rates calculated by fitting a constrained diffusion model to the initial 4 seconds of particle tracking data (Fig.2A-D). As has previously been shown using FRAP (5), the marker protein paGFP-LTI6b displays a significantly greater diffusion rate (D=0.063±0.003 μm^2^/s, p<0.01, Fig.2C & S2) when compared to the other proteins. The aquaporin PIP2A-paGFP (D=0.026±0.004 μm^2^/s) displays an enhanced diffusion rate when compared to FLS2-GFP (D=0.005±0.004 μm^2^/s, p<0.01) and PIN3-GFP (0.012±0.001 μ m^2^/sec, p<0.01, Fig.2C). The FLS2-GFP diffusion rate was significantly lower than that of PIN3-GFP (p≤0.05). Fitting a pure diffusion model to the first two points of each curve shows the same pattern for protein diffusion rates, demonstrating that our conclusions are robust to the choice of model although the precise diffusion values are different (Fig.2&S2). However, unlike the constrained diffusion rate for the proteins investigated, the constrained area occupied by the diffusing particle was shown to be the same for PIP2A-paGFP, FLS2-GFP and PIN3-GFP, with only paGFP-LTI6b showing a statistically significant increase in constrained area size compared to the other proteins (p<0.05-0.01, Fig.2D). Thus, we have demonstrated by single particle imaging that PM proteins move at different speeds within the membrane even when the areas that they move within are relatively similar in size.

**Figure 2:**
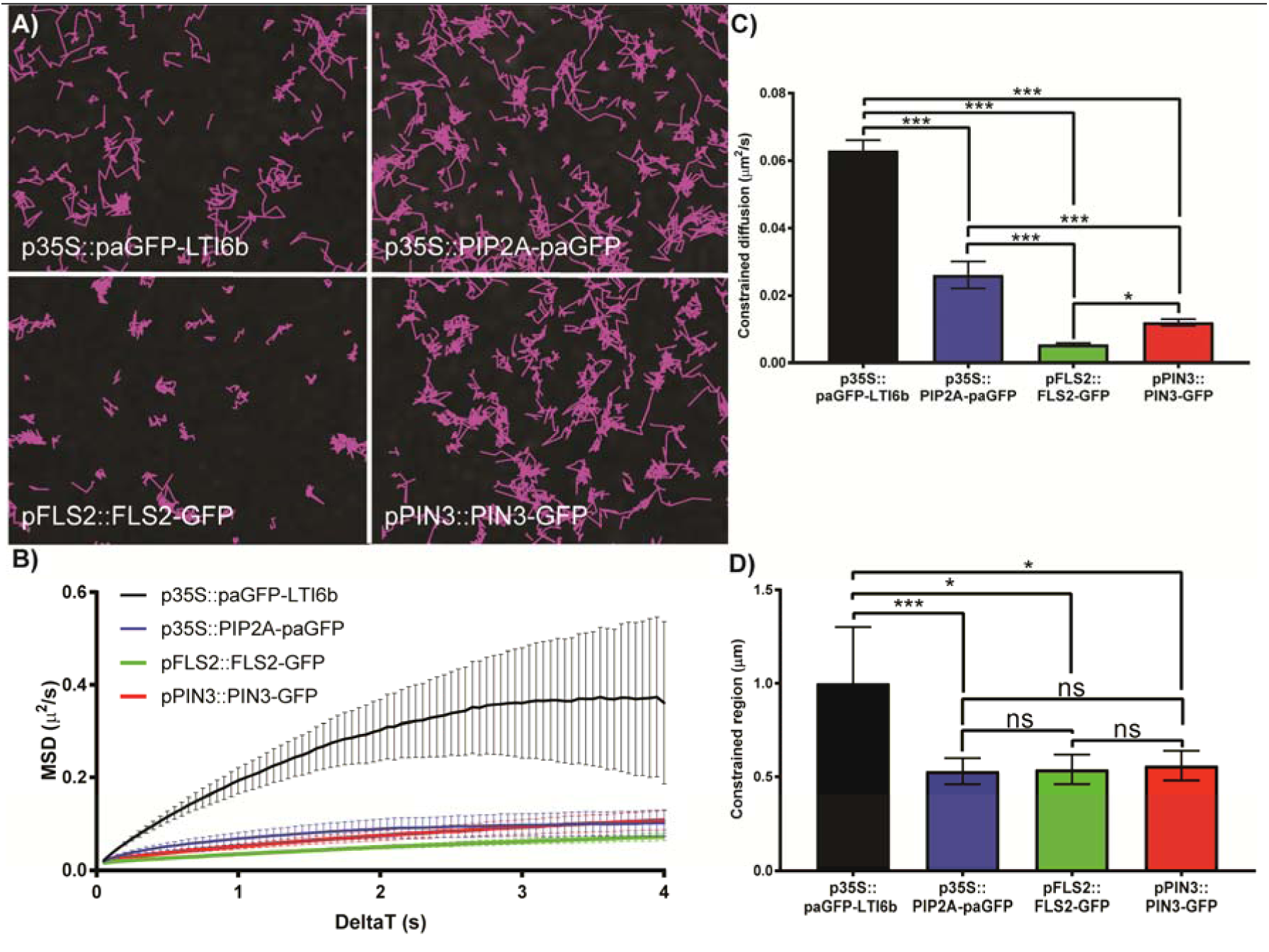
TIRF single particle tracking of PM proteins. A) TIRF-SPT of PM proteins in the hypocotyl membrane. Images show tracks followed by single labelled particles over 60s. Some proteins, e.g. FLS2-GFP are much more constrained in their lateral mobility than others. B) Mean Square Displacement curves. Curves that fall below a straight line corresponding to the initial gradient (as they all do) represent constrained diffusive movement. Error bars indicate bootstrap-estimated standard deviation. C) Constrained diffusion rate (μm^2^/sec) of proteins in the membrane. All proteins tested differ. D) Constrained region area (μm) proteins occupy in the membrane. *= p<0.05, ***= p<0.01, ns = not significant.

### The actin and microtubule cytoskeletons differentially regulate PM protein dynamics

The cell surface exists as a continuum containing the cell wall, PM and cytoskeleton (13). Previously it had been shown by FRAP that incubation of seedlings with cytochalasin D or oryzalin which depolymerize actin microfilaments or microtubules, respectively, did not affect the dynamics of ‘minimal’ membrane proteins (5). Here, upon actin or microtubule depolymerisation, no changes were observed in the constrained diffusion rate for PIN3-GFP and paGFP-LTI6b (Fig 3A&E, Supplemental movie 2). Interestingly, both showed a significant increase in constrained area size after actin depolymerisation (p<0.05, Fig 3B&F). Conversely, upon actin or microtubule depolymerisation, FLS2-GFP displayed an increase in protein diffusion rate, (Mock; D = 0.0053 ± 0.0004μ m^2^/s, Lat-B; D = 0.011 ± 0.002μ m^2^/s, oryzalin; D = 0.013 ± 0.002 μ m^2^/s, p<0.001, Fig 3C, Supplemental movie 2) but not in constrained area (Fig. 3D). This was also observed for instantaneous diffusion rates (Fig.S3). Therefore, the actin and microtubule cytoskeletons can differentially regulate the mobility of proteins in the membrane.

**Figure 3:**
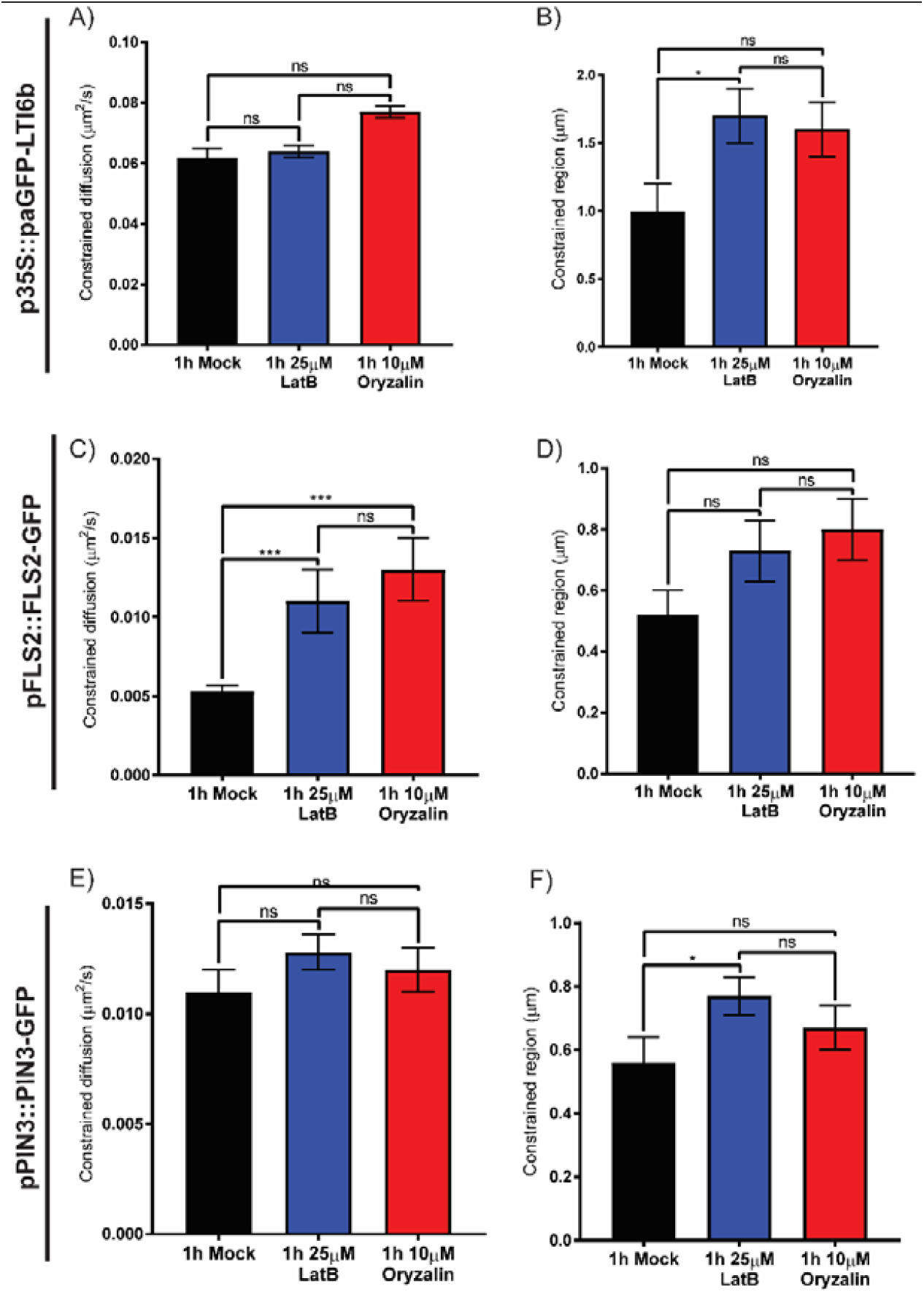
Actin cytoskeleton regulates the mobility of FLS2-GFP in the membrane. Plots show constrained diffusion rate (A, C, and E) and constrained area (B, D, and F) of single particles within the PM of hypocotyl epidermal cells in controls and after treatment with latrunculin B (LatB) and oryzalin to depolymerize the actin and microtubule cytoskeletons, respectively. A-B) p35S::paGFP-LTI6b, C-D) pFLS2::FLS2-GFP, and E-F) pPIN3::PIN3-GFP.FLS2-GFP becomes significantly more dynamic when either cytoskeleton is depolymerized. *= p<0.05, ***= p<0.01.

### The cell wall regulates PM diffusion rate, constrained area and nanocluster size

Lateral diffusion describes protein dynamics within the plane of a membrane. Previously it was shown using a combination of plasmolysis and protoplasting treatments that, upon removal of the cell wall constraint, protein lateral diffusion of ‘minimal’ PM proteins with extracellular-facing GFP is increased (5). Therefore, we hypothesized that the cell wall constrains the lateral diffusion rate of biologically functional proteins within the membrane. Here, we performed TIRF-SP imaging of paGFP-LTI6b, PIN3-GFP and FLS2-GFP in combination with pharmacological perturbation of the cell wall (Fig.4-5). PIN3-GFP and FLS2-GFP are both biologically active proteins with divergent function and were observed under control of their own promoters. The cellulose synthase specific herbicide DCB (24) and the pectin demethylesterase EGCG (25) were used to impair either cellulose synthesis or pectin methylation status (Fig.4-5) and hence the cell wall. Upon cell wall impairment with either, there was a non-significant trend towards increased constrained diffusion rate (Fig. 4B) and constrained area (Fig. 4C) for paGFP-LTI6b (Fig.4, Supplemental video 3). Therefore, over one hour of treatment with either drug, an alteration in cell wall structure did not dramatically alter paGFP-LTI6b dynamics within the membrane. There was however a significant increase in the instantaneous diffusion rate of paGFP-LTI6b upon cellulose or pectin perturbation of the cell wall (Fig.S5A&B, control; instantaneous D = 0.066 ± 0.005μ m^2^/s, DCB; instantaneous D = 0.085 ± 0.004 μ m^2^/s, EGCG; instantaneous D = 0.085 ± 0.003 μ m^2^/s). In addition, upon plasmolysis with either NaCl or mannitol, the paGFP-LTI6b diffusion rate was significantly increased in the PM (Fig.S4A-E, Supplemental video 4). Therefore, minor cell wall perturbation by impairing individual components does not affect the constrained diffusion rate of paGFP-LTI6b, but significant separation of the cell wall from the cell cortex and PM by plasmolysis does.

**Figure 4:**
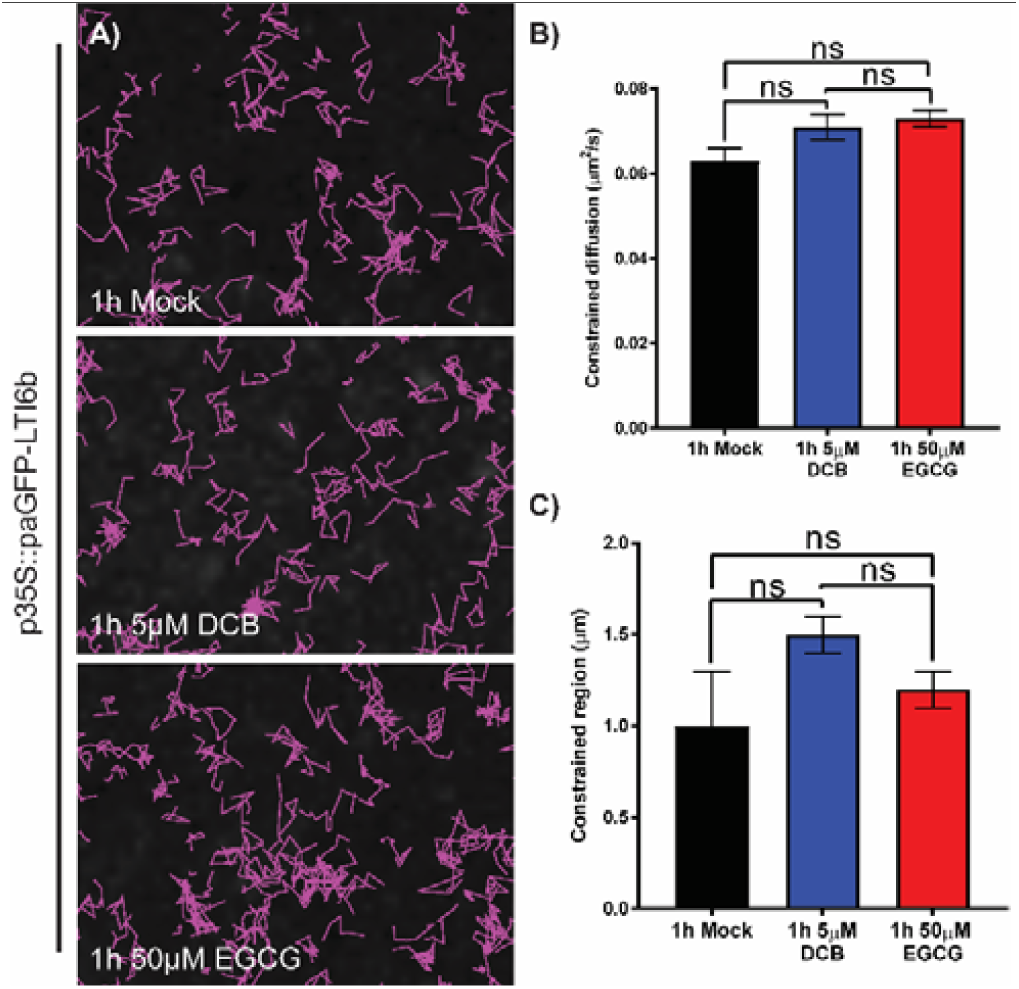
Single particle tracking reveals little effect of cell wall perturbation on paGFP-LTI6b dynamics. DCB was used to perturb cellulose synthesis and EGCG was used to perturb pectin methylation status of hypocotyl epidermal cells. A) p35S::paGFP-LTI6b in control, and after 5μM DCB and 50μM EGCG treatments for one hour each. Particles tracked over 60s. B) Constrained diffusion rate (μm^2^/sec) of proteins in the membrane tracked over 4 seconds. C) Constrained region (μm) proteins occupy in the membrane during 4 seconds. ns = not significant.

We also performed TIRF-SPT of the PM proteins PIN3-GFP and FLS2-GFP after cell wall perturbation (Fig. 5, supplemental movie 5). We chose PIN3-GFP and FLS2-GFP as their diffusion rates in untreated cells were reduced compared to paGFP-LTI6b and PIP2A-paGFP (Fig.2). In addition, PIN3 is functionally active in the hypocotyl as the flow of auxin is constant throughout plant development. Conversely, FLS2 should not be signalling in the absence of its ligand flg22 (26). In this study we tracked both active and non-active biologically functioning proteins and any similarities observed should demonstrate overall effects of the cell wall on PM protein dynamics. Unlike paGFP-LTI6b, both PIN3-GFP and FLS2-GFP showed significantly increased constrained diffusion rate and area upon treatment with either DCB or EGCG (Fig 5A-H). FLS2 diffusion was D = 0.0054 ± 0.0004μ m^2^/s in control, DCB; D = 0.0091 ± 0.001μm^2^/s, and EGCG; D = 0.013 ± 0.001μ m^2^/s, p<0.001. PIN3 diffusion was D = 0.012 ± 0.001μ m^2^/s in control, DCB; D = 0.0159 ± 0.0008μ m^2^/s, and EGCG; D = 0.018 ± 0.001μm^2^/s (p<0.05). Therefore, perturbation of either cellulose or pectin components of the cell wall results in these proteins diffusing faster and over a larger area (Fig 5). Furthermore, as a control, plasmolysis with either NaCl or mannitol and subsequent separation of the cell wall and PM caused an increase in diffusion rate and constrained area for both (Fig.S4F-O, Supplemental movie 6), with the exception of the constrained region for FLS2-GFP (Fig.S4J).

**Figure 5:**
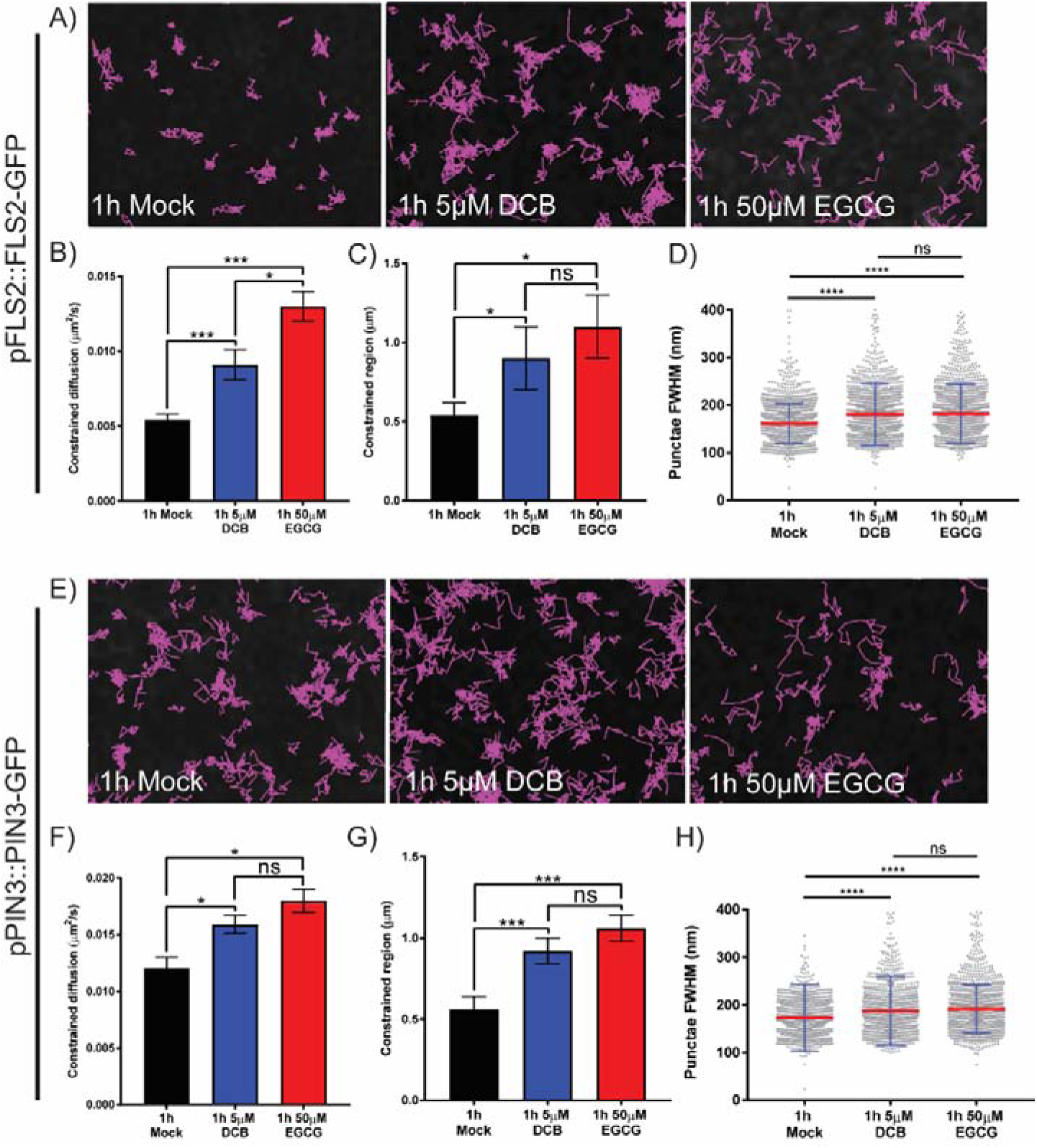
Cell wall perturbation alters diffusion rate, constrained area and cluster size of FLS2-GFP and PIN3-GFP. DCB was used to perturb cellulose synthesis and EGCG was used to perturb pectin methylation status of hypocotyl epidermal cells. A-D) Nanodomain characteristics of pFLS2::FLS2-GFP, and E-H) pPIN3::PIN3-GFP in either controls, or after treatment with 5μM DCB or 50μM EGCG for one hour. A and E) Track length of single particles over 60s. B and F) Constrained diffusion rate over 4s. C and G) Constrained region over 4s. D and H)FWHM measurement of cluster diameter. Box-and-whiskers plots, red line indicates mean, blue error bars represent standard deviation. There was a significant increase or trend towards increase in all nanodomain characteristics for both proteins after cell wall perturbation. *= p<0.05, **= p<0.01, ****=p<0.0001.

In combination with TIRF-SPT, Airyscan imaging of PIN3-GFP and FLS2-GFP demonstrated that nanodomain size significantly increases upon perturbation of either cellulose synthesis or pectin status (Fig 5D&H). FLS2-GFP control nanodomain size was FWHM = 161.4 ± 41.5nm, DCB; FWHM = 180.7 ± 65.35nm, and EGCG; FWHM = 182.1 ± 61.94nm (Fig. 5D). Nanodomain size after DCB and EGCG treatment was significantly greater than in controls (p≤0.0001, ANOVA), however there was no statistically significant difference between FLS2-GFP DCB and EGCG treated nanodomain size (p≥0.05, ANOVA). PIN3 control nanodomain size was FWHM = 173.1 ± 70.1nm, DCB; FWHM = 187.6 ± 72.29nm, and EGCG; FWHM = 191.5 ± 50.92nm (Fig. 5H). As with FLS2-GFP, PIN3-GFP nanodomain size was significantly greater after treatment with DCB or EGCG (p≤0.0001, ANOVA), however there was no significant difference between DCB and EGCG treated nanodomain size (p≥0.05, ANOVA).

Therefore, for FLS2-GFP and PIN3-GFP upon either plasmolysis, or cellulose and pectin disruption, there is an increase in constrained diffusion rate, constrained area, and nanodomain size. This demonstrates that the cell wall has a direct role in regulating both PIN3-GFP and FLS2-GFP protein dynamics and nanodomain size in the membrane.

### Discussion

#### Proteins reside in different sized nanodomains and display different dynamics in the plasma membrane

Here we have shown that several proteins form nanodomains within the plasma membrane which can be resolved with sub diffraction-limited imaging. Furthermore, the proteins we chose to image have diverse biological functions and have not been shown to have domains anchored into the cell wall as does, for example, FORMIN1 (18), AGP4 (5) or WAK1&2 (27). The auxin efflux transporter PIN2 has been shown using STED microscopy to form nanodomains in the membrane of between 100-200nm in diameter which is the same observed by us for PIN3-GFP using Airyscan imaging (Fig. 1 and reference (11). However in the same investigation BRI1 was found to have weak protein heterogeneity and hence nanodomain formation (11), which is in contradiction to our findings (Fig.S1) and those of others (28). We have imaged hypocotyl epidermal cells while the BRI1 study was conducted using root epidermal cells. Tissue-specific differences such as the cell wall status, which we and others have shown to be important for nanodomain size (Fig 5 and reference (5)), might explain these contradictory observations. We have shown that nanodomain size is significantly different for the various proteins investigated (Figs. 1&S1). Recent work has demonstrated that both FLS2 and BRI1 form nanodomains in the membrane (12, 28–30), which supports our study. However, the reported size for BRI1-GFP and FLS2-GFP nanodomains is significantly larger than we observe here (12). This is likely due to the imaging mode used and the image analysis methods employed.

Using TIRF Single Particle Tracking (TIRF-SPT), we have demonstrated that FLS2 and PIN3 have different diffusion rates within the plane of the PM. Furthermore, the dynamics of the proteins investigated are complex and not uniform. As shown previously, the paGFP-LTI6b diffusion rate is high relative to most other proteins thus far investigated (5). However it only has two amino acid residues projecting into the extracellular space compared to FLS2-GFP and PIN3-GFP which have larger extracellular domains (31, 32). ‘Minimal’ membrane proteins which are PM anchored and have an intracellular GFP tag have faster diffusion rates than ‘minimal’ membrane proteins which have extracellular GFP (2, 5). Therefore, with regard to investigation of PM protein dynamics, the study of functional biologically relevant proteins which contain extracellular domains is more instructive than marker proteins such as paGFP-LTI6b although the dynamics of biologically functional PM localised proteins which have no extracellular domains still needs to be investigated.

Protein domain diffusion rate heterogeneity exists in the plant PM for all proteins investigated in this study. This is similar to observations using dSTORM super resolution imaging of individual TCR molecules in activated human T cells (33) and proteins located in membrane sheets imaged with STED (34). Therefore, heterogeneity of membrane protein diffusion rates is a common theme across kingdoms. It is interesting to note that all proteins imaged also form differently sized nanodomains within the PM (Fig. 1 & S1). Heterogeneity of protein domain size and diffusion rate suggests that nanodomains of PM localised proteins must show substantial crowding / overlap within the membrane. However, we have only imaged one labelled nanodomain at a time in this study. It will be interesting to extend this work to investigate protein species heterogeneity within the imaged nanodomains. Protein association within nanodomains would convey rapid functionality in multi-protein response pathways. Additionally, it could account for how signalling pathways which rely on common components such as FLS2 and BRI1 can lead to environmental or development responses as has been shown previously (12). This could also account for cross talk between different pathways when components are localised to specific but partially overlapping nanodomains.

#### The actin and microtubule cytoskeleton can regulate the diffusion of FLS2 but not PIN3 and LTI6b

We have demonstrated that the actin and microtubule cytoskeletons do not uniformly regulate the dynamics of PM proteins. The actin and microtubule cytoskeletons only regulate the constrained diffusion rate of FLS2-GFP, which has increased lateral dynamics after depolymerization of either network (Fig. 3C). Both PIN3-GFP and paGFP-LIT6b showed no difference in diffusion rate upon cytoskeleton depolymerization, but did show an increase in the constrained area size when viewed as single particles (Fig. 3A-B & E-F). However, the constrained area was not altered for FLS2-GFP by cytoskeleton depolymerization (Fig. 3). PIP2A has been shown previously by sptPALM imaging to have an increased diffusion rate upon depolymerization of the actin cytoskeleton but no difference was reported for PIP2A upon oryzalin treatment to depolymerize the microtubule cytoskeleton (35). The actin and microtubule cytoskeleton regulation of some PM localised proteins is further demonstrated by a recent report showing that the pathogen perception signalling protein BIK1 co-localizes to microtubules but not the actin cytoskeleton (12). In addition, actin and microtubule depolymerisation resulted in loss of, and enlargement of nanodomain size of REM1.2, respectively (36). Furthermore, depolymerisation of the actin, but not the microtubule cytoskeleton reduces nanodomain density of LYK3 (10). However, it has also been demonstrated that for HIR1, microtubules govern nanodomain dynamics within the PM, preferentially to actin microfilaments (37). Differential regulation of proteins by the cytoskeleton would contribute to proteins forming differently sized nanodomains and having differing diffusion rates in the membrane, which we and others have observed. All proteins investigated in this study show differently sized nanodomains with different dynamics in the membrane (Fig. 1 & S1). The regulation of PM proteins by the cortical actin cytoskeleton has been investigated widely in mammalian cell systems and modelling has demonstrated that the actin cytoskeleton is sufficient to regulate heterogeneities in PM protein organisation (38). This could partly account for the differences we observe in PM nanodomains size and dynamics *in planta.*

#### The cell wall regulates PM nanodomain size and dynamics

To determine any effect that perturbations in different cell wall matrix components might have on the diffusion rate of proteins within the PM we perturbed cellulose synthesis and pectin methylation status. Neither of these treatments had a statistically significant effect on the constrained diffusion rate or area of paGFP-LTI6b in the membrane (Fig. 3). paGFP-LTI6b is an extremely mobile protein and shows very different characteristics during TIRF-SPT when compared to the biologically functioning PM proteins investigated. We hypothesize that due to the relatively fast diffusion rate of the protein in the PM and only having two residues in the apoplast, it is under relatively little constraint from the cell wall and hence, a minor cell wall perturbation over a short period such as those performed here with DCB and EGCG would not dramatically alter its diffusion rate. However, a major separation of the cell wall and PM during plasmolysis did significantly increase its diffusion rate in the membrane (Fig. S4).

PIN3-GFP and FLS2-GFP showed rapid changes in both constrained diffusion rate and constrained area upon cellulose or pectin disruption (Fig.5). Therefore the cell wall acts to constrain the lateral mobility of these proteins within the PM. We have demonstrated that cell wall structure also regulates nanodomain size (Fig. 5D&H). This is surprising as after cell wall perturbation for 20 minutes the cellulose synthase complexes are removed from the PM (15) but no other changes have been reported until much later with transcriptional changes, phytohormone induction and lignin deposition occurring at 4-7 hours of treatment (39). Therefore, minor cell wall perturbations rapidly affect PM nanodomain structure and dynamics. That such a short treatment has a profound effect on PM protein dynamics demonstrates how intimately related the cell wall and PM are. This could be an as yet undescribed mechanism of the plant cell that allows it to rapidly respond to mechanical stimuli. In addition, it is interesting that separating the cell wall and PM as occurs during plasmolysis results in increased diffusion of paGFP-LTI6b, whereas specifically impairing a single component over a short time frame did not. This could be because the cell wall has a global effect on the dynamics of all proteins with the severity depending on the size of any extracellular domains or residues. In addition, a subset of proteins with extracellular residues such as PIN3-GFP and FLS2-GFP might chemically interact with cell wall domains as has been demonstrated for Formin1 (5), and breakage of these chemical bonds resulting from plasmolysis might destabilize the entire membrane structure. The dense extracellular matrix of brain synapses has been shown to regulate the lateral mobility of AMSP-type glutamate receptors (40). Therefore, the role of extracellular matrices in governing the dynamics of PM proteins is common across kingdoms.

It would be interesting to determine if changes in nanodomain size affect the signalling functions of either PIN3 or FLS2 and subsequent hormone transport or ligand binding. Here we show using native promoter expression of tagged proteins that their dynamics and nanodomain size are regulated by the cell wall. The pathogen receptor protein FLS2 has lowered lateral mobility when treated with flg22 in protoplasts (41). Recently, it has been shown that flg22 treatment results in decreased dynamics of FLS2 nanodomains (12), confirming the FRAP result reported previously (41). This has also been demonstrated for the aquaporin PIP2A which, upon salt stress, co-localizes with the membrane nanodomain marker FLOT1 and shows changes in its mobility within the membrane (42). Additionally, LYK3, upon ligand binding and host cell infection shows reduced dynamics and increased stability in the membrane (10). In addition, membrane nanodomains have been shown to be important for the activation of receptor-mediated signalling upon ligand perception and subsequent clathrin-mediated endocytosis (28). Therefore, given that the cell wall plays a role in regulating the size of these nanodomains and their dynamics, cell wall regulation of PM nanodomains is of fundamental importance to signalling *in planta.*

To conclude, we have shown that a number of PM proteins form nanodomains within the PM and that these are of sufficient size for imaging using sub-diffraction limited techniques such as the Zeiss Airyscan system. These nanodomains are of different sizes and their dynamics and size can be differentially regulated by the actin and microtubule cytoskeletons and the cell wall. As yet, very limited information exists as to how PM proteins form nanodomains. We demonstrate here that the cell wall plays a key role in regulation of protein nanodomain size and lateral mobility for the pathogen receptor FLS2 and the auxin transporter PIN3. We hypothesize that the cytoskeleton and cell wall slow nanodomain dynamics sufficiently to allow relatively static distribution of functional proteins so that they are well placed spatially for optimum association.

### Materials and Methods

#### Plant material

The *Arabidopsis thaliana* lines used have been previously described; p35S::paGFP-LTI6b (5), pFLS2::FLS2-GFP (26), pPIN3::PIN3-GFP (43), p35S::PIP2A-GFP (44), p35S::PIP2A-paGFP (45), pREM1.3::YFP-REM1.3 (6) and pBRI1::BRI1-GFP (46). Seeds were surface sterilized in 70% ethanol for 5 minutes, 50% bleach for 5 minutes and washed four times with water. Seeds were placed on square agar plates composed of ½ strength MS with MES and 0.8% Phytagel. Seedlings were then stratified on plates for 2 days at 4°C in the dark and then placed into a growth chamber set to 16:8h long days, 23°C, and 120μ-Einstein’s light intensity for 5 days before imaging.

#### Chemical treatments

*A. thaliana* seedlings were treated in 8ml dH_2_O in 6 well plates for 1 hour with the following concentrations, all made from 1000X stocks; 5μM DCB, 50μM EGCG, 0.5M mannitol, 100mM NaCl, 2.5μM latrunculin-B and 10μM oryzalin. DCB, isoxaben, latrunculin-B and oryzalin were dissolved in DMSO and EGCG was dissolved in ethanol.

#### Confocal microscopy

Seedlings were imaged after five days of growth by mounting them in dH_2_0 on microscope slides with no1.5 coverslips. Slides and coverslips were held down with micropore tape. A Zeiss LSM880 equipped with an Airyscan detector was used. Airyscan imaging was performed using 488 and 514nm excitation for GFP and YFP respectively. Lasers were used at 1% transmission with a dual 495-550BP and 570nm long pass filter. For standard confocal imaging the same emission wavelength was imaged with a GaAsP detector. To avoid chlorophyll autofluorescence a 615nm short pass filter was used. A 100x/1.46 DICM27 Elyra oil immersion lens was used for all imaging. A 5X zoom was used to image flat membrane sheets and imaging conditions were all set according to Zeiss optimal Airyscan frame size (for 5X zoom, 404×404). Frame sizes were kept the same for standard confocal imaging. For single particle experiments, sample size (n) = a minimum of 12 cells imaged across 3 biological replicates per condition, the number of single particles tracked per condition is displayed in Table S1. For all Airyscan data n = ≤64 nanodomains were measured / cell for 36 cells across three biological repeats, exact numbers for each condition can be seen in Table S2.

#### Airyscan image analysis

PM protein nanodomain size was determined by imaging using the above conditions. Using the FIJI implementation of imageJ, an 8X8 grid was placed over the image and line profiles determined for the brightest nanodomain in each grid cell. The full width half maximum (FWHM) of these line profiles was then determined and this data was collated in Graphpad Prism version 7. Scatter dot plots were produced with error bars denoting the standard deviation. ANOVA with multiple comparisons was used to assess nanodomain size differences for different proteins. Kymographs were produced from 55 subsequent images comprising 8 seconds of imaging the PM. The Multiline kymograph in FIJI was used to produce a kymograph with the line originating in the bottom left corner at a 45 degree angle to the top right for each data-set.

#### TIRF-SP Imaging

TIRF imaging was performed as described in (5) using an inverted microscope (Axio Observer, Zeiss) equipped with a 100X objective (α-Plan-Apochromat, NA = 1.46; Zeiss) and TIRF slider (Zeiss), 488-nm laser excitation (Stradus Versalase, Vortran), HQ525/50-nm emission filter (Chroma), and an electron-multiplication CCD (iXon+; Andor). The exposure time was 50 ms.

#### TIRF-SPT - particle tracking

From single particle tracks, mean squared displacement (MSD) curves were calculated as MSD(*ΔT*)=<|**r_i_**(*T+ΔT*)-**r_i_**(*T*)|^2^> where |**r_i_**(*T+ΔT*)-**r_i_**(*T*)| is the displacement between position of track *i* at time *T* and time *T*+*ΔT* and the average is over all pairs of points separated by *ΔT* in each track. The errors in the MSD curve were calculated by repeating the MSD curve calculation 200 times, each time on a different synthetic dataset created by randomly resampling with replacement the tracks present within each dataset, and the datasets present (bootstrap resampling (47)). The distribution of MSDboot_*j*_(*ΔT*) curves about the MSD curve for the resampled data, MSD(*ΔT*), should be close to the distribution of MSD(*ΔT*) about the true MSD curve (47). Therefore a posterior sample of 200 MSD curves MSDpost_*j*_ (*ΔT*) can be calculated from these 200 bootstrap MSD curves MSDboot_*j*_ (*ΔT*) (*j*=1.200).

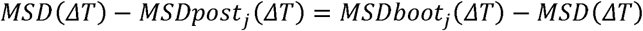

So

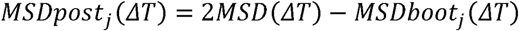

Subsequent model fits (see below) were performed on each posterior MSD curve sample to naturally yield joint posterior samples of the fitted model parameters suitable for determining confidence intervals, error bars and statistical tests. A*χ*^2^ fit was performed for each posterior sample using the standard deviation of the posterior MSDs at each *ΔT* as the error estimate for calculating *χ*^2^.

The models fitted were free diffusion with parameters diffusion rate *D* and localisation error σ_*loc*,_ which was fitted to the first two points on the curve, for which

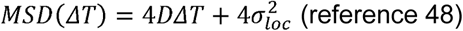

and constrained diffusion with parameters initial diffusion rate *D*, confinement region size *L* and localisation error σ_*loc*_ where

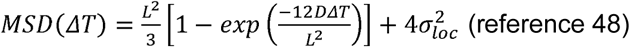

The confidence intervals for each parameter were chosen as the midpoint ± half width of shortest interval containing 69% of the posterior probability for that parameter.

We assume for the null hypothesis that posterior samples 1 and 2 correspond to the same value of quantity *x*, the probability of a given difference*Δ x* is the same as the measured probability of *Δ x* about its mean, i.e.

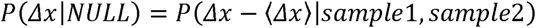

The probability that | *Δ x* | is at least;⟨ *Δx* ⟩) given the null hypothesis is then

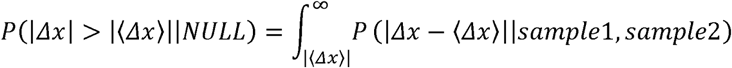

We used this as a non-parametric p-value for the null hypothesis that the two posterior samples measure the same value. In the case of normally distributed posteriors from normally distributed sample measurements this gives the same p-values as the 2-sided Welch’s t-test.

## Supporting information

## Acknowledgements

JFM was funded by BBSRC grant BB/K009370/1 awarded to JR. AFT was funded by an Oxford Brookes Nigel Groome studentship. DR, SEDW, SB and MM are funded by the STFC. Access to the STFC funded facilities for TIRF-SPT imaging was provided by programme access grant 16130041 to JR and CH. We thank T. Ott, S. Robatzek and J. Chan for generously providing seed lines. We would also like to thank all members of the plant cell biology section at Oxford Brookes for insightful discussions.

## List of supplemental Materials

1. **Supplementary Table 1** Number of tracks analysed per construct per treatment for single particle imaging.

2. **Supplementary Table 2** Number of nanodomain size measurements per construct per treatment for Airyscan imaging.

3. **Supplemental Figure 1** Comparison of Confocal and Airyscan imaging of PM nanodomains.

4. **Supplemental Figure 2** Instantaneous diffusion values for TIRF single particle tracking of PM proteins.

5. **Supplemental Figure 3** Instantaneous diffusion values for TIRF single particle tracking of p35S:: paGFP-LTI6b, pPIN3::PIN3-GFP and pFLS2::FLS2-GFP during cytoskeleton perturbation.

6. **Supplemental Figure 4** Plasmolysis causes changes in single particle dynamics for p35S::paGFP-LTI6b, pFLS2::FLS2-GFP and pPIN3::PIN3-GFP.

7. **Supplemental Figure 5** Instantaneous diffusion values for p35S::paGFP-LTI6b, pPIN3::PIN3-GFP and pFLS2::FLS2-GFP during cell wall perturbation.

8. **Supplemental Movie 1** TIRF single particle tracking of p35S::paGFP-LTI6b, p35S::PIP2A-paGFP, pFLS2::FLS2-GFP and pPIN3::PIN3-GFP shows they diffuse at different rates and occupy differing sized areas within the PM.

9. **Supplemental Movie 2** TIRF single particle tracking of p35S::paGFP-LTI6B, p35S::PIP2A-paGFP, pFLS2::FLS2-GFP and pPIN3::PIN3-GFP during control, actin (Lat-B) and microtubule (Oryzalin) depolymerisation.

10. **Supplemental Movie 3** TIRF single particle tracking of p35S::paGFP-LTI6b in the PM during cell wall perturbation.

11. **Supplemental Movie 4** TIRF single particle tracking of p35S::paGFP-LTI6b in the PM during plasmolysis.

12. **Supplemental Movie 5** TIRF single particle tracking of pFLS2::FLS2-GFP and pPIN3::PIN3-GFP during cell wall perturbation.

13. **Supplemental Movie 6** TIRF Single particle tracking of pFLS2::FLS2-GFP and pPIN3::PIN3-GFP during plasmolysis.

