## Supplementary material for "The cell wall regulates dynamics and size of plasma-membrane nanodomains in *Arabidopsis*"

**Supplemental Figures.**

**Supplemental Table 1. Number of tracks analysed per construct per treatment for single particle imaging.**

|  | **p35S::paGFP-LTI6B** | **pFLS2::FLS2-GFP** | **pPIN3::PIN3-GFP** | **p35S::PIP2A-paGFP** |
| --- | --- | --- | --- | --- |
| **Mock** | 2799 | 2617 | 5059 | 8500 |
| **5µM DCB** | 2173 | 2315 | 5749 | N/A |
| **50µM EGCG** | 3692 | 2724 | 5562 | N/A |
| **20µM Isoxaben** | 4819 | 2117 | 7214 | N/A |
| **100mM NaCl** | 2130 | 2006 | 2055 | N/A |
| **0.5M Mannitol** | 1249 | 1364 | 2078 | N/A |
| **2.5µM Latrunclin B** | 5062 | 2781 | 4959 | N/A |
| **10µM Oryzalin** | 2325 | 3400 | 4183 | N/A |

**Supplemental Table 2. Number of nanodomain size measurements per construct per treatment for Airyscan imaging.** Nanodomain comparison data used in figures 1 and S1. Cell wall perturbation used in Figure 5.

| **Genotype** | **Condition** | **Experiment** | **# measured** |
| --- | --- | --- | --- |
| pBRI1::BRI1-GFP | Mock | nanodomain comparison | 2301 |
| pFLS2::FLS2-GFP | Mock | nanodomain comparison | 2245 |
| pPIN3::PIN3-GFP | Mock | nanodomain comparison | 2235 |
| p35S::PIP2A-GFP | Mock | nanodomain comparison | 1308 |
| pREM1.3::YFP-REM1.3 | Mock | nanodomain comparison | 1420 |
| pPIN3::PIN3-GFP | Mock | Cell wall perturbation | 1564 |
| pPIN3::PIN3-GFP | 5µM DCB | Cell wall perturbation | 1234 |
| pPIN3::PIN3-GFP | 50µM EGCG | Cell wall perturbation | 1470 |
| pFLS2::FLS2-GFP | Mock | Cell wall perturbation | 2164 |
| pFLS2::FLS2-GFP | 5µM DCB | Cell wall perturbation | 2249 |
| pFLS2::FLS2-GFP | 50µM EGCG | Cell wall perturbation | 2207 |

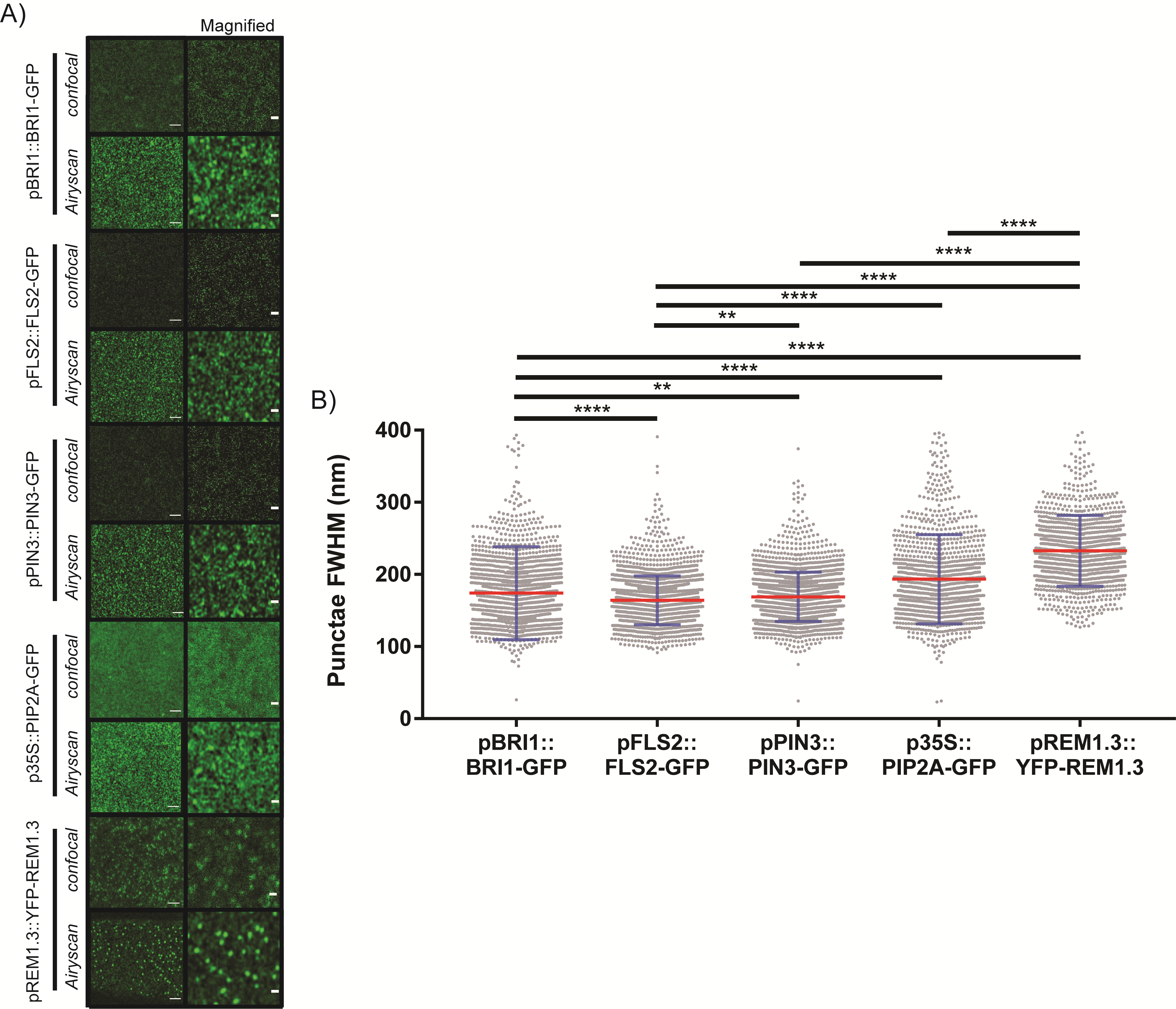

**Supplemental Figure 1.** **Comparison of Confocal and Airyscan imaging of PM nanodomains**. A) Comparison of Confocal and Airyscan imaging of a number of PM localised proteins, scale bar denotes 500nm. B) Scatter plot showing nanodomain size calculated using FHWM of line profiles. FLS2-GFP, PIN3-GFP and YFP-REM1.3 also shown in Fig. 1. ** = p≤ 0.01, **** = p≤ 0.0001 ANOVA. Red line denotes mean, blue error bars denote standard deviation. Please note, FLS2, PIN3 and REM1.3 data also in figure 1 but shown here for complete comparison. Number of nanodomain quantified = BRI1; 2301, FLS2; 2245, PIN3; 2235, PIP2A; 1308, REM1.3; 1420 collected from 3 experimental repeats.

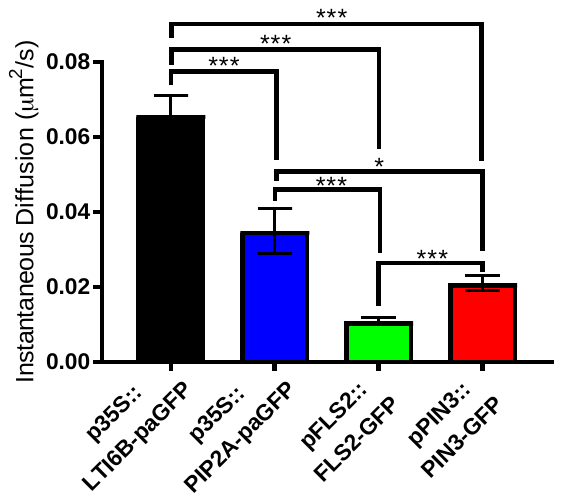

**Supplemental Figure 2.** **Instantaneous diffusion values for TIRF single particle tracking of PM proteins.** Instantaneous diffusion rates of p35S::paGFP-LTI6b, p35S::PIP2A-paGFP, pFLS2::FLS2-GFP and pPIN3::PIN3-GFP PM localised proteins determined by TIRF-SPT. *=p<0.05, ***=p<0.01.

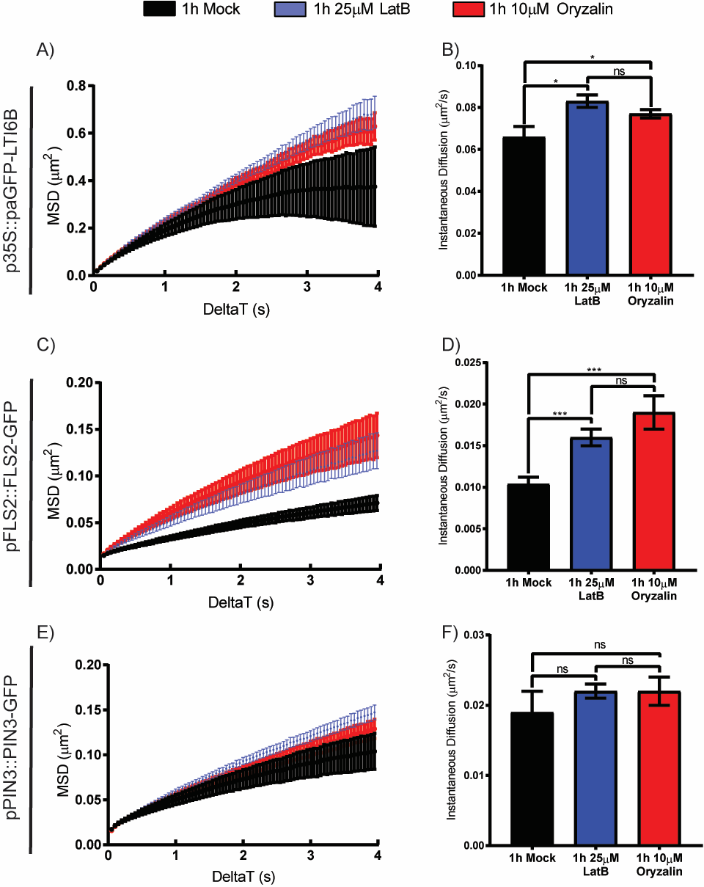

**Supplemental Figure 3.** **Instantaneous diffusion values for TIRF single particle tracking of p35S:: paGFP-LTI6b, pPIN3::PIN3-GFP and pFLS2::FLS2-GFP during cytoskeleton perturbation.** Instantaneous diffusion rates determined by TIRF-SPT during actin (Lat-B) and microtubule (oryzalin) cytoskeleton perturbation. *** = p<0.01, *=p<0.05, ns=not significant.

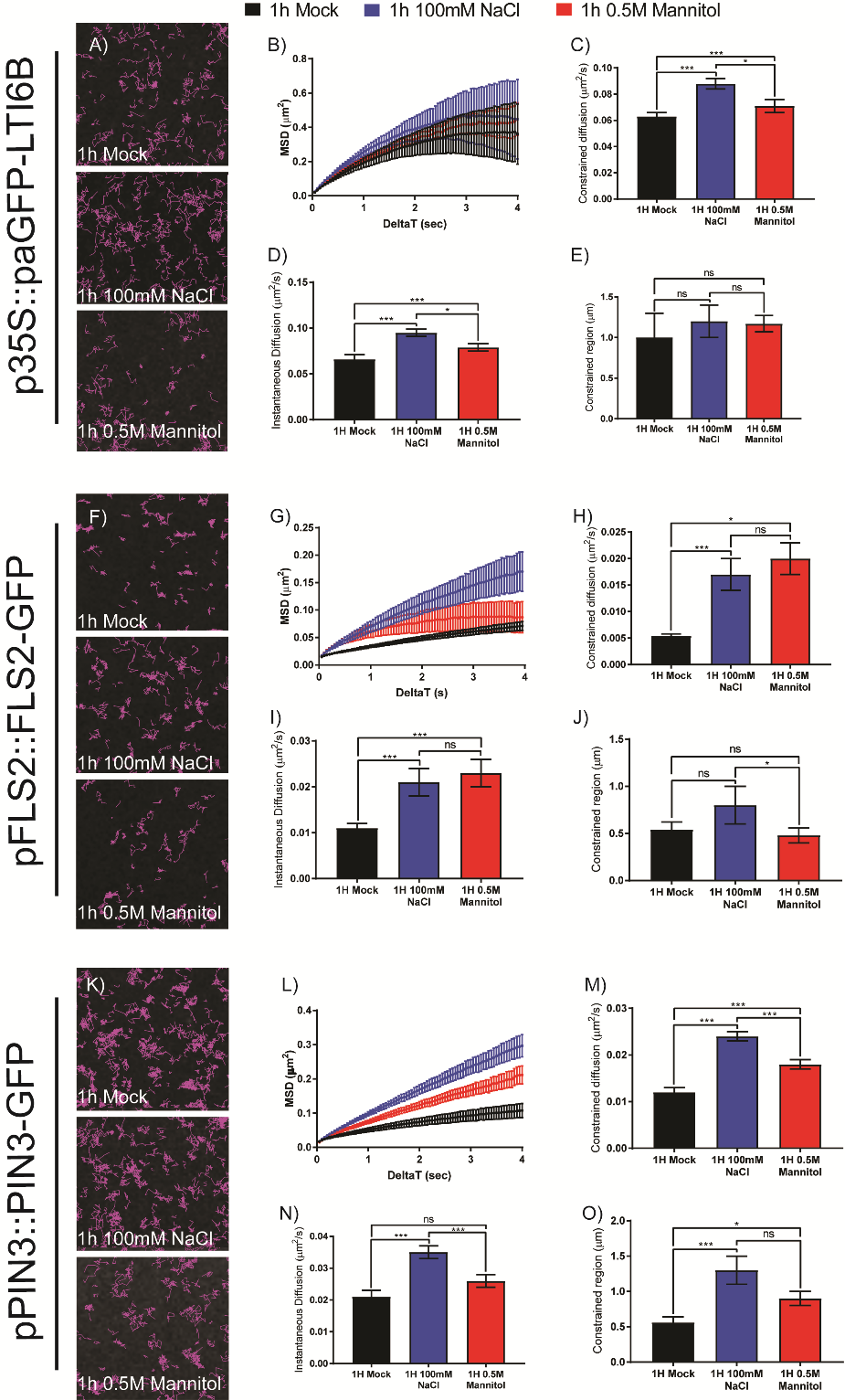

**Supplemental Figure 4.** **Plasmolysis causes changes in single particle dynamics for p35S::paGFP-LTI6b, pFLS2::FLS2-GFP and pPIN3::PIN3-GFP.** A) p35S::paGFP-LTI6B single particle tracks in mock, 100mM NaCl and 0.5M Mannitol plasmolysis. B) MSD curve of p35S::paGFP-LTI6B plasmolysis treatments. C) Instantaneous diffusion rates of p35S::paGFP-LTI6B single particles tracked during plasmolysis treatments. D) Constrained diffusion rates of p35S::paGFP-LTI6B particles tracked over 4s during plasmolysis treatments. E) Constrained area of p35S::paGFP-LTI6B particles tracked over 4 seconds during plasmolysis treatments. F) pFLS2::FLS2-GFP single particle tracks in mock, 100mM NaCl and 0.5M Mannitol plasmolysis. G) MSD curve of pFLS2::FLS2-GFP plasmolysis treatments. H) Instantaneous diffusion rates of pFLS2::FLS2-GFP single particles tracked during plasmolysis treatments. I) Constrained diffusion rates of pFLS2::FLS2-GFP particles tracked over 4 seconds during plasmolysis treatments. J) Constrained area of pFLS2::FLS2-GFP particles tracked over 4 seconds during plasmolysis treatments. K) pPIN3::PIN3-GFP single particle tracks in mock, 100mM NaCl and 0.5M Mannitol plasmolysis. L) MSD curve of pPIN3::PIN3-GFP plasmolysis treatments. M) Instantaneous diffusion rates of pPIN3::PIN3-GFP single particles tracked during plasmolysis treatments. N) Constrained diffusion rates of pPIN3::PIN3-GFP particles tracked over 4 seconds during plasmolysis treatments. O) Constrained area of pPIN3::PIN3-GFP particles tracked over 4 seconds during plasmolysis treatments. ns=not significant, *=p<0.05, *** p<0.01.

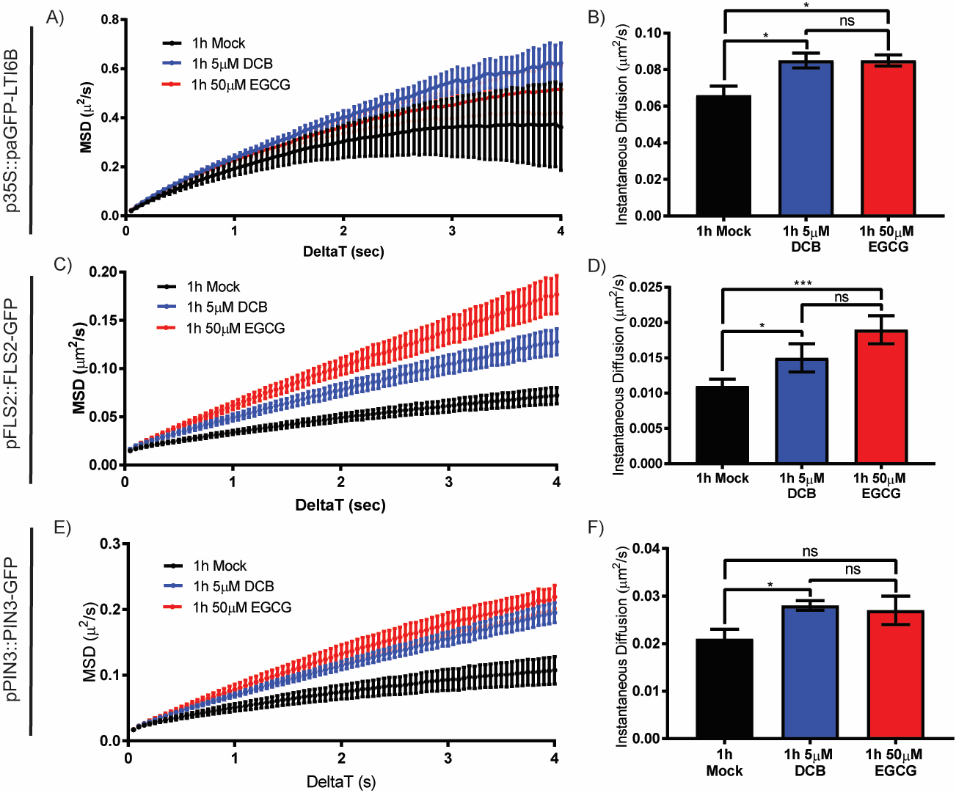

**Supplemental Figure 5.** **Instantaneous diffusion values for p35S::paGFP-LTI6b, pPIN3::PIN3-GFP and pFLS2::FLS2-GFP during cell wall perturbation.** Instantaneous diffusion rates of p35S::paGFP-LTI6B, pFLS2::FLS2-GFP and pPIN3::PIN3-GFP with mock, 5µM DCB and 50µM EGCG treatment. ns=not significant, *=p<0.05.
